# Isolation and characterization of toxic Cyanobacterial Communities distribution in Lake Tana, Amhara Regional state, Ethiopia

**DOI:** 10.1101/659748

**Authors:** Adugnaw Admas, Aklilue Agida, Smegnew Melese, Amare Genetu

## Abstract

Lake Tana is the second largest lake next to Victoria Lake in Africa. This Lake is exsposed to agriculture run off, industrial and Urban waste since it has not buffer zone to protect any invading materials to the water body. The accumulation and growth of blue-green algae in water body call public attention because of its health concern. Some cyanobacterial species produce toxin microcystins and anatoxin molecules can affect the liver, kideney and nervous system of animals because the toxic molecules of released from some toxic cyanobacteria may lead the organs to fail the metabolice system. The major exposure pathways of these toxins are through drinking water and recreations including swimming. In addition, these problems happen through food web. Those cyanobacteria’a are spread and growth in diffrent water body via agriculture run off, industrial and urban wastes and other nutrient source of N and P. The main goal of this scientific study was to assess the distribution of potential toxic cyanobacteria in Lake Tana. Cyano bacterial distributions were studied in different water bodies of Lake Tana. From the water bodies that have visible algal blooms and not observed algal blooms were investigated. Identification of cyano bacteria was conducted using microscope by following appropriate procedures. In the present studied area Anabaena sp, Nostoc sp., Chlorella sp., and Microcystis aeruginosa sp. were dominantly exist. Therefore, Maybe the numbers of fishe depleted in Lake Tana due to the presence of microcystine molcule source, those are Anabaena sp.,Nostoc sp. and Microcystis aeruginosa sp.

## 1. Introduction

Lake Tana is found in East Africa, North West Ethiopia. The Lake is surrounded by lagoons and wetlands and has 40 tributaries (rivers and streams), on which Gilgel Abay, Ribb, Gumara and Megech account for 93 % of the total inflow (Setegn *et al*. 2008). The lake was formed 20 million years ago by a lava extrusion that functions as a natural dam (UNESCO, 2011).

Lake Tana water shade consists of 347 kebeles and 21 districts in four administrative zone of Gondar and Gojam (IFAD, 2007). The total area of catchment is 15,319 square kilometer (Conway, 2000). In the Northern Part of the Lake, the most notable landmarks are existing in it’s edge. It is far 60 kilometers from Gondar, before Gondar it was the capital city of Ethiopia and the name so called is Goregora peninsula. In this peninsula different monastery like Mandaba, Debresina and other attractive historical monasteries are existed (Cheesman .R.E., 1935).

It is the largest fresh water source in Ethiopia and has a surface area of 3,156 sq km, a minimum depth of 9 meter and maximum depth of 14 meter length of 84 km and width of 68 km (Amare, 2011).

Lake Tana basin is one of the three sections of Eastern Afromontane biodiversity hotspot (Mittermeier *et al.*, 2004) is a center of biodiversity and endemism which contains 65 fish species with about 70 % of the fish species in the lake being endemic (Goshu *et al.*, 2010). The faunal diversity also comprises of 277 mammals, 862 birds, 201 reptiles and 63 amphibians with some endemic to Ethiopia. Ethiopia as a country has the fifth largest flora in tropical Africa estimated that 6,500-7,000 species of higher plants, of which 12 % are endemic and some are found in the Lake Tana basin. The agro biodiversity comprises of many field crops, indigenous medicinal plants and church forest islands of biodiversity (EBI, 2009) and over 113 woody plant species identified in Lake Tana (Alelign, 2007).

The basin (Lake Tana Basin and Nile Valley) are the repositories of ancient indigenous culture, linguistics, history and ancient civilization. Thousands years ago, science and technology in Medicine, Mathematics, Agriculture, Architecture, communication or writing system, commerce, etc.) Were practiced in the Nile basin and it is assumed that the current world’s technology is originated from the Nile basin and African rift valley (UNEP, 2006; Wikipedia).

Lake Tana has 37 islands of which 20 have Ethiopian Orthodox churches and monasteries with an enormous historical and cultural value, dating back up to the 14th century (Beckham *et a*l., 1954).

Lake Tana is the growth and development collider for Ethiopia, South Sudan, Sudan and Egypt. It is the main sources of the Blue Nile, which contributes about two-thirds of the water volume of the Nile in which the country’s electric power and agricultural sectors are greatly, depend on. For instance, besides irrigation, the fish resource potential of Lake Tana is 50,000 metric tons per year, and it supports an estimated population of over (Goshu *et al*., 2010; Wikipedia). Overall, this lake and the adjacent wetlands provide directly or indirectly a livelihood for more than 500,000 people (Vijverberg *et al*., 2009).

The lake also balances the climatic condition of the region. The northern pick (i.e. Mount Dashen) and Lake Tana is the green belt which prevents the spread of the Sahara desert and offer wetter climates. The climate of the Eastern Afromontane Hotspot has been relatively constant over recent geological history due to high levels of biodiversity, endemism and thermo – coolant factors of the lakes and mountains (Balmford *et al*., 2001; Borgess *et al*., 2007).

The Lake also supports hydroelectric powers. In this regard, the lake plays a role to control environmental pollution. Hydropower’s are most importantly clean energy sources with relatively negligible production of noxious gases or solid/ liquid wastes, and therefore the dam can largely prevent environmental pollution, climate change, global warming and related public health consequences. Levels of greenhouse gases (GHG) emission from hydro-power are relatively low (Kaunda *et al*., 2012). The life cycle GHG emission factors for hydro-power technologies are around 15–25 g CO2 equivalent per kWhel. These are very much less than those of fossil-fuel power generation technologies which typically range between 600–1200 gram CO2 equivalents per kWhel (Lenzen, 2008). Power generated from129 million tons of coal increases pollutants to the atmosphere, including 7.7 million tons of particulates and 296 million tons of CO2 (Mantunge *et al*., 2015)

Even though, Lake Tana and its surrounding wetlands are of immense ecological value and provide the means of existence for millions of people, however, it is threatened by multifaced problems associated with unwise human activities in the catchment areas and invasive species affects their natural features. Overall, degradation of catchment areas, drainage for agriculture, overgrazing and unsustainable resource use, water hyacinth, siltation and pollution from agrochemicals and urban sewerages are the major threats significantly affecting Lake Tana ecosystems (Mequanent and Sisay, 2015; Kindie, 2001). Anthropogenic pollution from fecal pollution and agricultural sources as fertilizers, insecticides and herbicides are also detected in the lake especially in the shores and river mouths (Goshu *et al.*, 2010).

Recent studies show a serious decline in fish stocks due to the spread of the aquatic weed water hyacinth around fish spawning grounds (Melesse and Samuel, 2018). In Addition, Because of agriculture run off, industry waste and other N and P source is directly discharged to the Lake Tana, this has caused the increased number of algal blooms in Lake Tana, which lead to reduce the number of fishes in the lake (Amhara National Regional State Water and Mines Resources Development Bureau reported, 1990).

Cyanobacteria, commonly known as blue-green algae, are bacteria that contain photosynthetic pigments similar to those found in algae and plants. Their ability to fix nitrogen directly from the atmosphere gives them a competitive advantage over other algae. Blue-green algae blooms typically occur when plant nutrients such as phosphorus and nitrogen are in plentiful supply. However, factors needed for a bloom are complex.

Blue-greens cannot maintain an abnormally high population for long and will rapidly die and disappear after 1-2 weeks. If conditions remain favorable, another bloom can replace the previous one, making it appears as one continuous bloom lasting for up to several months. Toxic blue-greens are an emerging public health issue (Stone and Bress, 2007). The primary exposure pathways of concern have been drinking water and recreational exposure. Consumption of fish containing blue-green toxins represents a poorly studied weather how much it is toxic or not (Ibelings and Chorus, 2007; Stone and Bress, 2007; Wilson *et al*., 2008). Microcystins are heat stable and do not break down during cooking (Harada *et al*., 1996). These compounds are suspected liver carcinogens, which could prove significant to humans following continuous, low-level exposure. Due to its instability, anatoxin-a is more difficult to analyze than microcystins. Several species of cyanobacteria can grow abundantly under favorable natural environmental conditions and form high biomass called water blooms which often is associated with eutrophication (Kanoshina *et al 2003*, Milan 2007). Cyanobacterial blooms commonly occur in many temperate lakes (Hudnell, 2008) and also in coastal areas (Vargas-Montero and Freer, 2004). The blooms may initially appear green and later turn blue-green, sometimes forming a ‘scum’ on the water surface. These blooms are considered a natural phenomenon, but in recent years their frequency has increased considerably (Kanoshina *et al* 2003, Carmichael 2008). Agricultural runoff and other effluents to fresh and marine water bodies and wetlands have resulted in increased nutrient enrichment of phosphorous and nitrogen (Kanoshina *et al* 2003), thus providing favorable conditions for the growth of toxic cyanobacteria.

Most of the harmful effects of cyanobacterial blooms have been reported from freshwater ecosystems. Several cyanobacteria blooms have also been reported from brackish and marine waters and may have harmful effects on humans and animals (Stewart, 2008). The bloom of a marine Cyanobacterium, *Trichode smiumerythraeum* causes sickness, dermatitis and other discomforts (Vargas-Montero and Freer, 2004).

Therefore, our hypothesis was that the fish may be reduced by the microcystin which is released from harmful cyanobacteria in Lake Tana.

Hence, our research attempted to assess the presence of the harmful cyanobacteria in Lake Tana by identifying the species using microscope.

## 2. MATERIALS AND METHODS

Water samples and cyanobacterial mats were collected from Lake Tana since October 2017 and Feb. 2018 in Goregora side at 1190SE12015’45’’ N37^0^18’11’’E, Grand Hotel side at 28^0^NE11^0^36’2’’N37^0^22’60’’E, St.Michael side at 280^0^11^0^36’41N37^0^22’33’’E and St.mariam at 79^0^E11^0^37’9’’N37^0^20’49’’E by plastic bottles and using 3ml /L Lugol’s solution for micro algae sample nutrient source till sample analysis in laboratory. Then, 250 ml of water sample was filtered through the filter membrane with pore size of 0.22 μm. After filtration, membrane was washed in 5 ml of autoclaved water. The filtered microalgae were growing in the standard BGI medium for 15 days. The culture media of BGI contained MgSO_4_.7H_2_O 0.4g; MgCl_2_.6H_2_O 0.7g; CaCl_2_.2H_2_O 0.5g; KH_2_PO_4_ 0.3g; K_2_HPO_4_ 0.3g; (NH_4_)_2_SO_4_ 0.5g; H_3_BO_3_ 0.26g; CuSo_4._ 5H_2_o 0.5g; MnCl_2_. 4H_2_o 0.5g; Mo 0.06g and ZnSo4. 7H_2_o 0.7g in 1 Liter of deionized water.

### 2.1 Isolation of blue-green algae

The cultivated micro algaes sample by BGI media were transferred to 2% solid agar media, then different uni culture were observed after 10 days in this media. Finally, using a microscope the cultured cyanobacteria were identified.

## 3. Result and Discussion

The cyanobacterial species *Anabaena sp., Microcystis aeruginosa sp.* and *chlorella* sp. were found in St.Michel Monastery side of Lake Tana, cyanobacteial species from Goregora side of the Lake Chlorella and *Nostoc sp.* were investigated. Three cyanobacterial species were found at St.Mariam side of Lake Tana those were *spirulina platensis, Microcystis aeruginosa sp.* and *chlorella sp*. Also, the dominant cyanobacteria species, *Anabaena sp*. were founded in the Grand hotel side.

**Figure 1.**
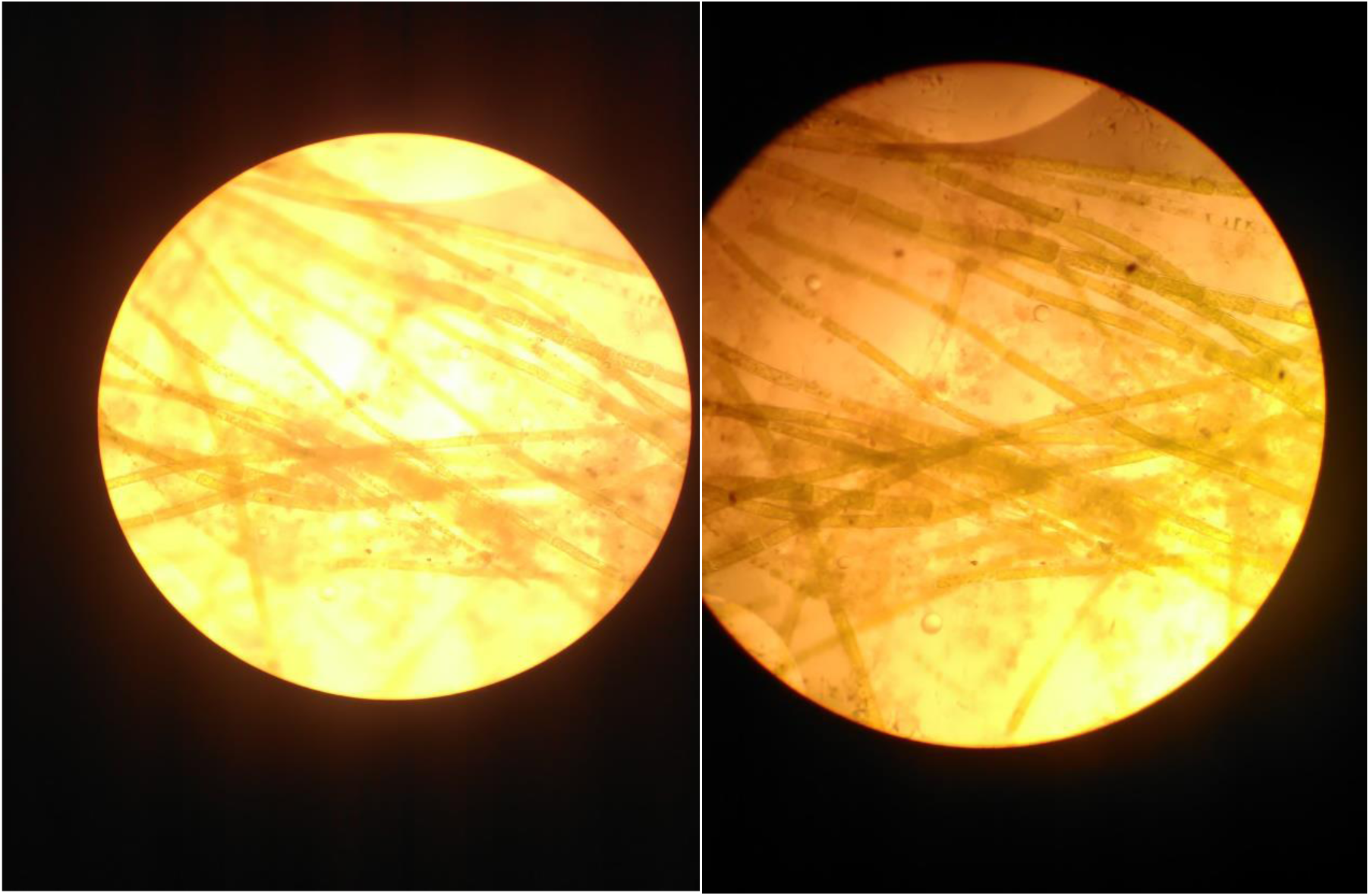
Grand hotel (*Anabaena species*)

**Figure 2.**
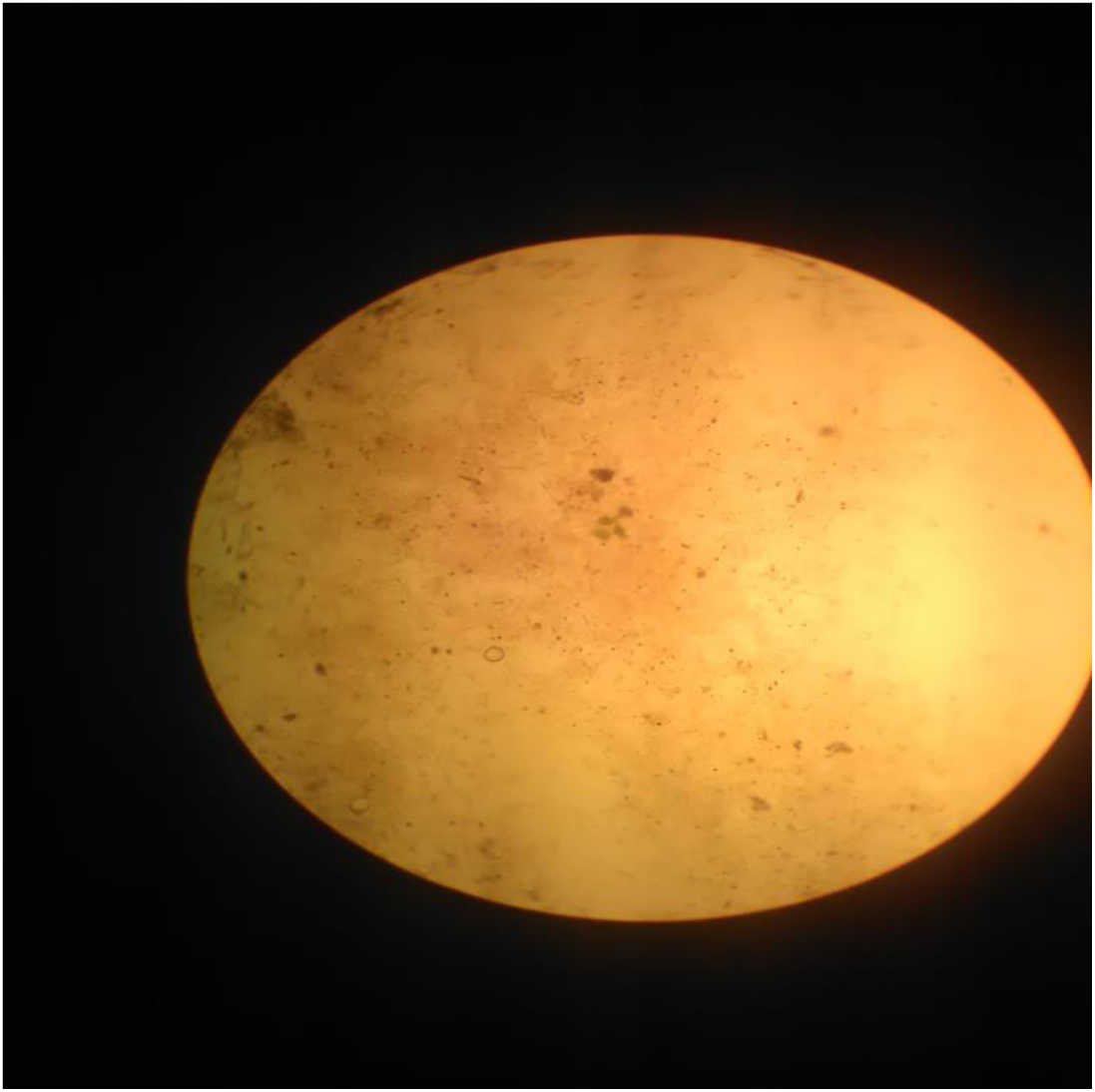
St.Michael (*Microcystis aeruginosa species*)

**Figure 3.**
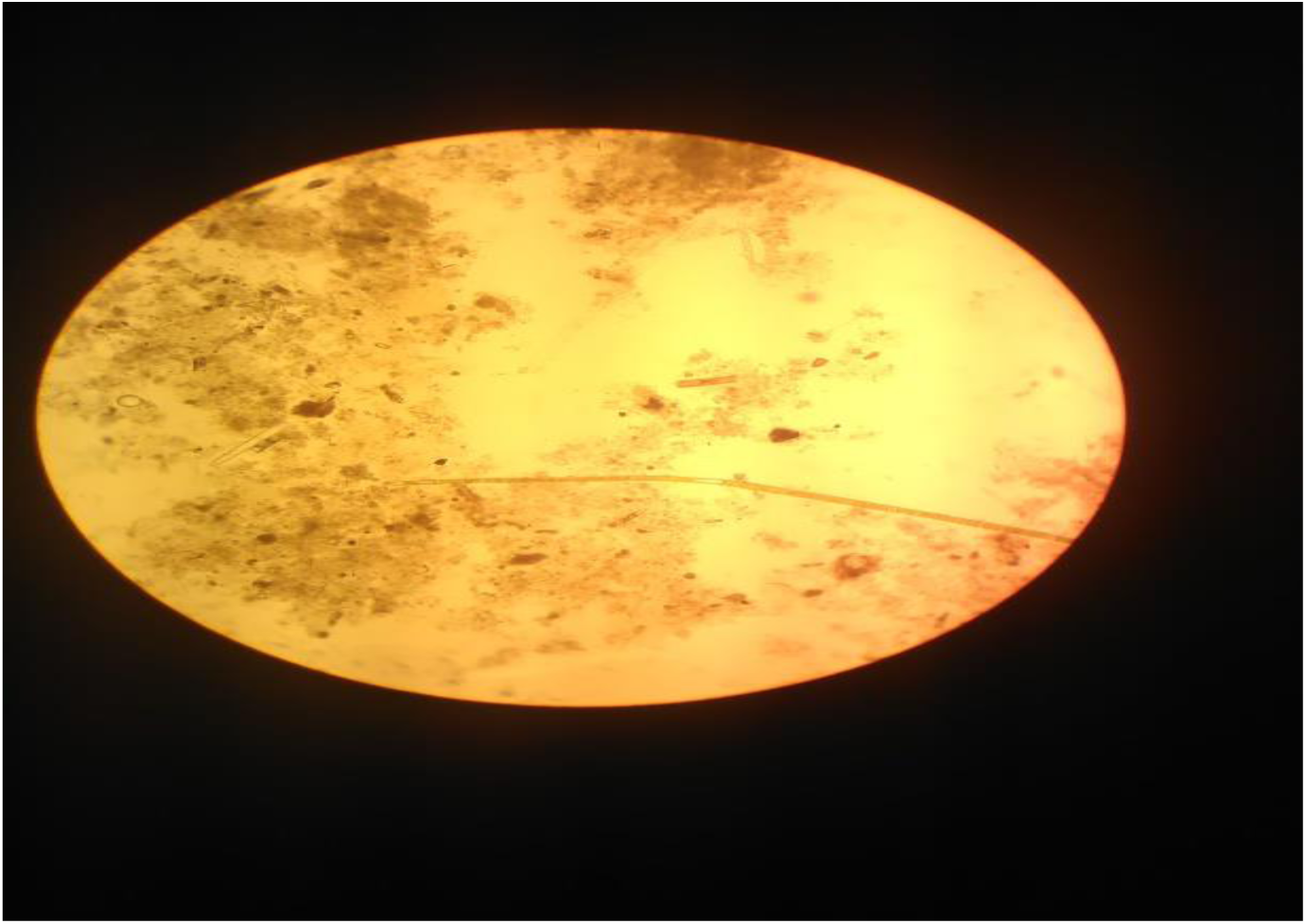
St.Mariam (*spirulina platensis, Microcystis aeruginosa sp.*and *chlorella sp.*)

**Figure 4.**
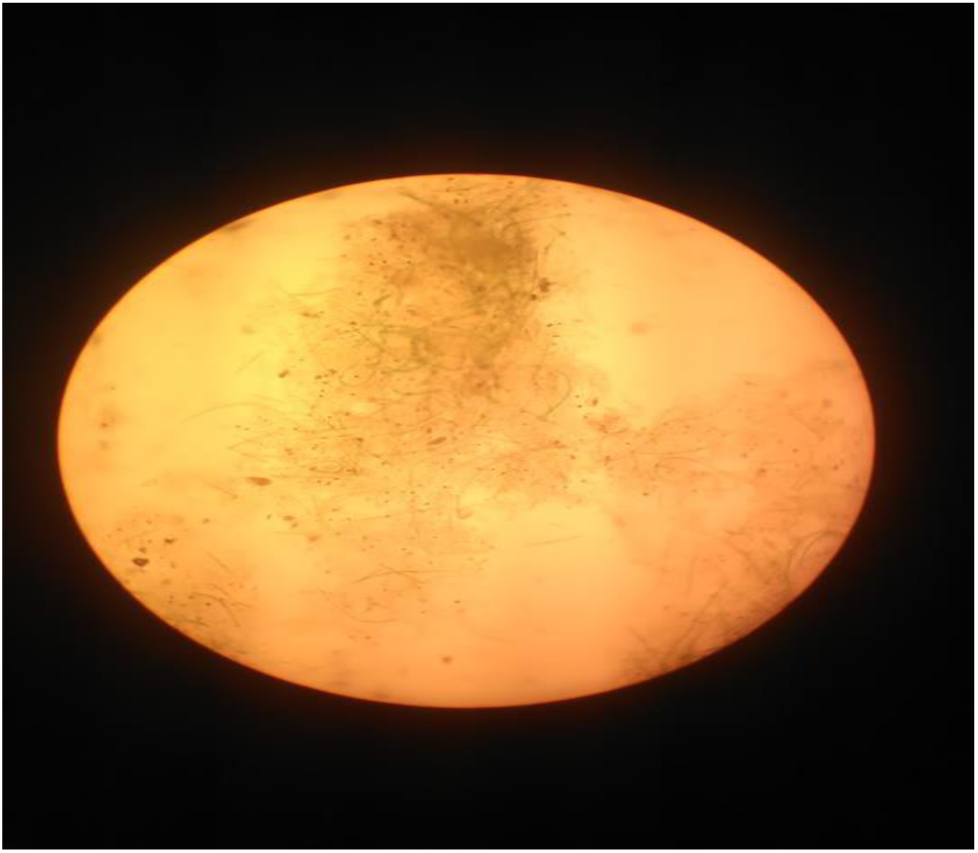
Goregora (*Nostoc sp.*,.and *chlorella sp.*)

Cyanobacteria are well known for their source of a multitude of highly toxic and/or allelopathic compounds. The toxic compounds include various cyclic peptides (the hepatotoxic microcystins) and alkaloids (the potent neurotoxins and the hepatotoxic cylindrospermopsin), which have been studied both from a toxicological and a biological perspective (Hudnell, 2008). In this study, *Microcystis aeruginosa sp., Nostoc sp.*, and *Anabaena* sp., were dominantly existing in Lake Tana. Those cyano bacteria (blue-green algae) are potential toxic and cytotoxic effects source of different animals, for example from the genus *Nostoc sp*., nostocyclamide, a cyclic hexapeptide inhibits growth in algae and bacteria (Jüttner *et al*. 2001, Becher *et al*. 2009). Nostocarboline that - functionally similar to anatoxin-a(s) - is an inhibitor of acetylcholinesterase and the first serine protease inhibitor of an alkaloid structure that has been described (Hirata *et al.* 1996). Nostocine inhibited the growth of various algae and cultured plants (Nagatsu *et al*. 1995; Gromov *et al*. 1991).

Microcystins are produced by *Anabaena, Fischerella, Gloeotrichia, Nodularia, Nostoc, Oscillatoria*, members of *Microcystis*, and *Planktothrix*. Microcystins are the most widespread cyanobacterial toxins and can bioaccumulate in common aquatic vertebrates and invertebrates such as fish, mussels, and zooplankton. Microcystins primarily affect the liver (hepatotoxin), but can also affect the kidney, and reproductive system (United States of America Environmental protection agency, 2008)

Most of the studied water bodies are under an excessive nutrient loading from agricultural and urban runoff, it leads the lake to become eutrophicated. In this eutrophicated area toxic microcystin protein producer Microcystis *aeruginosa sp*., *Anabaena and Nostoc sp*., from the collected water samples were found.

## 4. Conclusion and recommendation

The present results indicate that potentially toxic cyanobacteria that synthesis microcystine molecules were *Microcystis aeruginosa sp, Anabaena* sp., and *Nostoc sp*., were occurred in the studied areas of Lake Tana. Therefore, as we know these cyanobacteria are very toxic and harmful to aquatic organisms, so it may affect the fish in the lake. This research finding recommend LakaTana should have standard buffer zone to protect the Lake from agriculture runoff, urban, industrial waste and other water pollutant source since the nutrient source of cyanobacteria is from outside the Lake and finally, harmful algal blooms in Lake Tana should be continuously monitoring to protect human and animal health.

## 5. Acknowledgements

This work was supported by grants of Ethiopian Environment and Forest Research Institute.

